# Through the Random Forest: Ontogeny as a study system to connect prediction to explanation

**DOI:** 10.1101/2022.08.11.503703

**Authors:** Sophia Simon, Paul Glaum, Fernanda S. Valdovinos

## Abstract

As modeling tools and approaches become more advanced, ecological models are becoming more complex and must be investigated with novel methods of analysis. Machine learning approaches are a powerful toolset for exploring such complexity. While these approaches are powerful, results may suffer from well-known trade-offs between predictive and explanatory power. We employ an empirically rooted ontogenetically stage-structured consumer-resource model to investigate how machine learning can be used as a tool to root model analysis in mechanistic ecological principles. Applying random forest models to model output using simulation parameters as feature inputs, we extended established feature analysis into a simple graphical analysis. We used this graphical analysis to reduce model behavior to a linear function of three ecologically based mechanisms. From this model, we find that stability depends on the interaction between internal plant demographics that control the distribution of plant density across ontogenetic stages and the distribution of consumer pressure across ontogenetic stages. Predicted outcomes from these linear models rival accuracy achieved by our random forests, while explaining results as a function of ecological interactions.

## Introduction

As ecologists expand the range and depth of ecological theory they necessarily integrate advanced modeling techniques and computational tools to produce increasingly complex models. This increase in model complexity has been driven, in part, by the long-standing debate on the relationship between diversity and stability in ecosystems ^1^. Differing levels of diversity (in terms of number of species and interactions) directly affects the number of interacting variables and, therefore, model complexity ^2,3^. Indeed, numerous studies continue to raise unique sources of complexity and separate mechanisms tying diversity to increased stability. These mechanisms include weak species interactions ^4^, adaptive foraging ^5^; allometric scaling of interaction strength ^6^; and omnivorous ^7^, mutualistic ^8^, or high-order interactions ^9^. These diverse sources of model complexity inspired by ecological processes, their unique mechanistic effects on model dynamics, and the nonlinearities they produce emphasize ecologists’ need for an adaptable methodology that can be deployed across model frameworks. Our work shows how machine learning can be used to develop such an adaptable methodology while addressing its typically associated limits in interpretability.

Modern machine learning algorithms are flexible, powerful tools used to study many complex systems and are increasingly applied to ecological datasets ^10^. Recently, ecologists have applied machine learning algorithms to predicting species interactions from empirical trait data ^11^, improving estimation of viral host ranges from incomplete datasets ^12^, and examining food web responses to variable functional diversity across trophic levels ^13^. While these machine learning approaches are powerful, they have been called “black boxes” because their high predictive power does not guarantee highly interpretable results ^14^. Recently, methods for the productive interpretation of machine learning results have been reviewed in the hopes of transforming this “black box” into a “translucent box” ^10^. However, developing broadly interpretable results is difficult. This is partially because machine learning algorithms necessarily rely on direct model output, which can potentially limit ecological generalizability as model complexities lead to incommensurable (i.e., lacking common measurement standards) results across different formulations and parameterizations ^13^.

One key source of ecological complexity is demographic heterogeneity, frequently modeled as stage-structured organismal ontogeny ^15^. Explicitly modeled ontogenetic stages increase the number of unique ecological actors within a single species. This consequently increases the total number of species interactions in an ecosystem because each ontogenetic stage tends to have its unique interactions with other species or stages ^16^. Stage-structured models have made clear the importance of demographic heterogeneity in influencing population dynamics, particularly in plants ^17,18^. At the community level, the few food web studies addressing the role of organismal ontogeny on community stability have found mixed results. Rudolf & Lafferty ^19^ found that diet shifts across organismal development (ontogenetic niche shifts) destabilize food webs. However, in formulating a community model without ontogenetic niche shifts, de Roos ^20^ found that explicit ontogeny stabilizes food webs so long as ontogenetic development involves substantial asymmetries between juveniles and adults. These qualitatively different results indicate that the specific demographic and ecological formulation of ontogeny influences its effect on broader dynamics. Again, here we see the potential for incommensurable results.

Studying how dynamics change with specific ecological/ontogenetic structures is best addressed initially through simpler, more tractable models that facilitate a deeper sensitivity analysis across model formulations, providing a useful basis and reference as model complexity is scaled up to represent entire communities. Modeling plant-herbivore interactions is well suited for this purpose given that plants frequently have clear ontogenetic stages or size distinctions that affect their intra- and interspecific interactions, and plants’ autotrophic nature allows us to build small-scale tractable models that can form the basis of larger food webs. Furthermore, as a consequence of the long history of plant ontogeny in ecology, there are empirical resources to aid in vetting basic model formulation ^21^.

Therefore, we focus on a plant-herbivore model with empirically informed ontogenetic stage structure and parameterization. Using this simulation model’s parameters as random forest input features (see Methods), we then implement a machine learning based analysis of model behavior and show its utility in identifying and categorizing context dependent dynamics. Given the high degree of context-dependent results observed from our simulation model (and others; ^13^), we use this opportunity to not only study the dynamic contingencies across ontogenetic formulations, but also take steps towards developing broadly applicable mechanistic ecological understanding of seemingly incommensurable results while maintaining the high predictive accuracy of our random forests.

## Methods

### Model Development and Justification

We implemented the plant ontogeny framework from Glaum & Vandermeer (2021; see Eq. 1, Fig. 1, and Appendices S1.1 & S1.2). Plant ontogeny is divided into three stages: a seed bank (*S*_1_), non-reproductive seedlings (*S*_2_), and fecund adults (*F*). While there are clearly a variety of potential ontogenetic structures, a three-stage ontogenetic structure provides initial tractability in analysis and is well-represented across plant species (see Appendix S1.1). Public data repositories ^21^ hold nearly 1200 instances of plant species demographics represented by a three-stage structure representing 107 unique species. These three-stage plant species span a wide phylogenetic distribution, representing both eudicots and monocots across 3 phyla, 3 classes, 28 orders, 45 families, and 87 genera. Botanically, these species represent shrubs, succulents, trees, epiphytes, annuals, and herbaceous plants. Finally, three-stage plant species are also wide-spread geographically, representing eleven terrestrial ecoregions across all continents except Antarctica, making three-stage structures worthy of concentrated theoretical evaluation. For further details, please see SI File 1 and Appendix S1.

**Figure 1:**
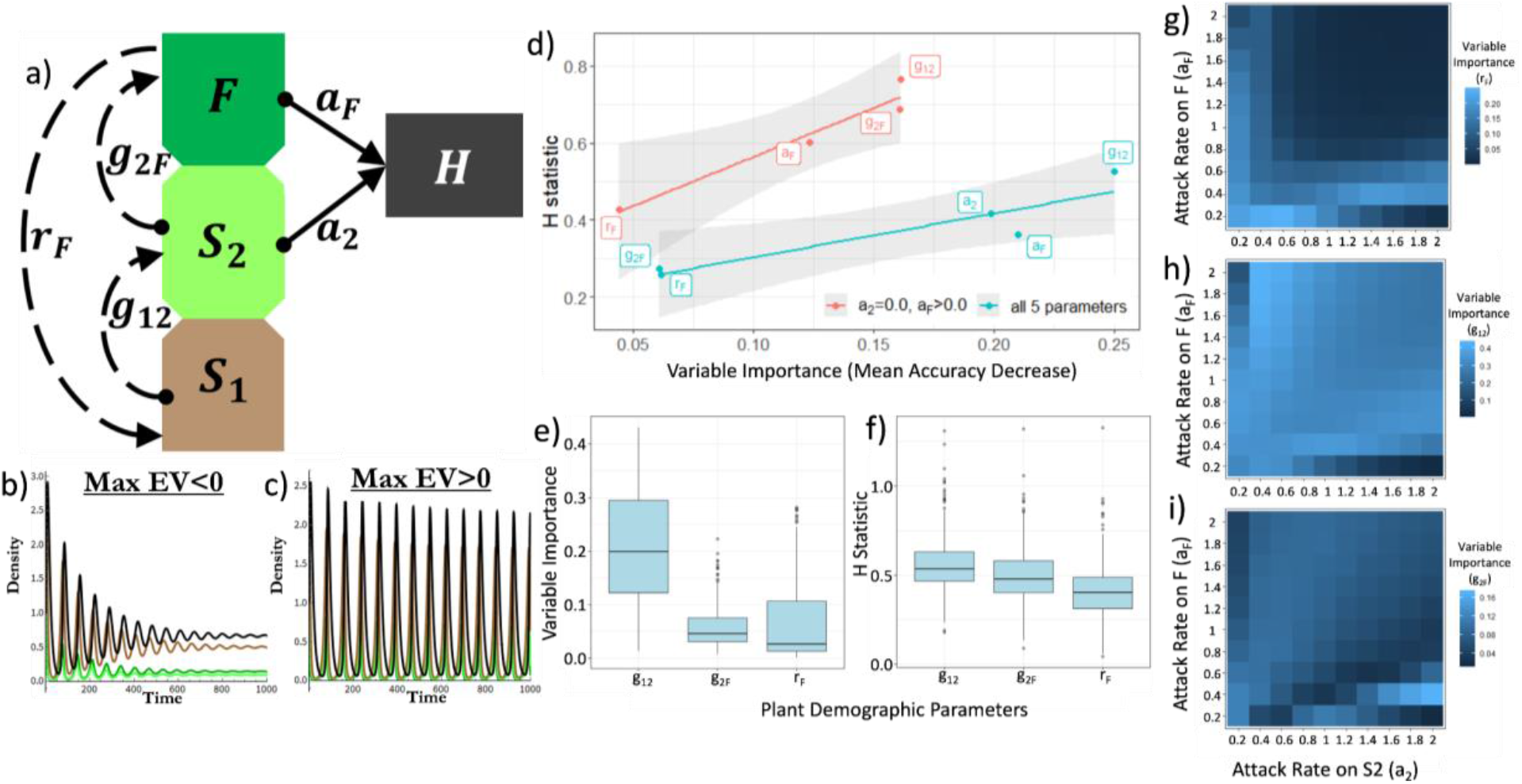
Model & Random Forest overview. a) Model diagram and major parameters mediating model flow. Dashed lines indicate plant population dynamics, solid lines indicate trophic interactions. Arrows indicate direction of density gain and circles indicate direction of density loss. Variables and parameters detailed in Methods. b) Example of stable population trajectories indicated by negative max eigenvalue. c) Example of persistent population oscillations indicated by positive max eigenvalue. Line colors for b) &c) correspond to Fig 1a. d) Relationship between Variable Importance and interactivity (H-statistic) of parameters in Random Forest output. Blue line: RF model across all five parameters, AUC:0.998 (α_*g*1_=α_*g*2_=α_*F*_=0.1; *h*_2_=*h*_*F*_=0.5). Red line: RF model with single stage herbivory (*a*_2_=0); AUC=0.984 (α_*g*1_=α_*g*2_=α_*F*_=0.1; *h*_2_=*h*_*F*_=1). Shaded regions represent standard error. e,f) Box and whisker plots detailing range of Variable Importance (e) and H-statistic (f) for each demographic rate in random forests run with set attack rates where *a*_2_ and *a*_*F*_ vary between 0.2 and 2.0 (α_*g*1_=α_*g*2_=α_*F*_=0.1; *h*_2_=*h*_*F*_=0.5). Boxes represent the interquartile range with the horizontal line showing the median, the lower box showing the 25 percentile, and the upper box showing the 75 percentile. Upper and lower lines extending from the boxes show the most extreme values within 1.5 times the 75th and 25th percentile respectively. Outliers are shown as single dots. g,h,i) Heatmaps showing changing importance for each demographic rate across different allocations of consumption across plant stages. Here we found consistently but only slightly better performance with mtry=2.

The stage-structured plant in our model is consumed by a single herbivore species (*H*) (Eq 1; Fig 1a). Real world herbivory can certainly involve multiple species interactions. However, specialization of herbivore species on a single plant taxa (especially at the family level) is common (especially amongst insect herbivores) and geographically widespread ^22,23^. The modeled herbivore is limited to eating the vegetative stages (*S*_2_ & *F*) as the role of “seed predator” is rarely filled by a species which also consumes vegetative tissue, again especially in insects due to the requirements of different mouth parts. Among herbivore species which consume vegetation (specialist or otherwise), past work has cataloged examples of herbivores exhibiting a range from clearly distinct to non-existent ontogenetic preferences in their plant resources ^24–28^. Still other examples have found variable preferences, potentially resulting from local plant chemistry, leaf palatability, and microclimate ^27,29^. Reflecting this empirical range, we vary the focus of herbivory to either specialize on a particular ontogenetic stage (adult or seedling) or both stages to varying degrees. The model is shown in Eq. 1 with each component (germination, consumption, etc.) labeled as different sub-functions. Sub-functions and model parameters are detailed in Table 1.

**Table 1:** Model parameters and sub-functions. Parameter descriptions include values used in numerical analysis. Color highlighted parameters indicate varied parameters during parameter sweeps. Green highlighted parameters indicate parameters whose value range was varied in factorial parameter sweeps (see Appendix S1.3) for analysis via random forest. Orange highlighted parameters were varied as part of sensitivity analysis. Note, for each simulation α_g1_=α_*g*2_ =α_*F*_. All parameter rates are per capita. Note, the 5 green highlighted parameters were used as input features in our random forests.

| Parameter | Definition |
| --- | --- |
| $r_F$ | Intrinsic reproduction (seed) rate of fecund plant individuals ($F$). $0.2 \leq r_F \leq 4.0$ |
| $g_{12}$ | Germination rate of seeds ($S_1$) into seedlings ($S_2$). $0.12 \leq g_{12} \leq 0.88$ |
| $g_{2F}$ | Maturation rate of seedlings ($S_2$) into fecund adults ($F$). $0.12 \leq g_{2F} \leq 0.88$ |
| $a_F$ | Attack rate of the herbivore ($H$) on the fecund adult plant population ($F$). $0.0 \leq a_F \leq 2.0$ |
| $a_2$ | Attack rate of the herbivore ($H$) on the seedling population ($S_2$). $0.0 \leq a_2 \leq 2.0$ |
| $c_{FH}, c_{2H}$ | Conversion rate of eaten adult plants ($c_{FH}$) or seedlings ($c_{2H}$) into herbivores. $c_{FH} = c_{2H} = 0.6$ |
| $\alpha_F, \alpha_{g1}, \alpha_{g2}, \alpha_{FS}$ | Strength of density dependence affecting seed production, seed germination, seedling maturation, and seed survival respectively. $\alpha_{FS} = 0.001$; $\alpha_{g1}, \alpha_{g2}, \alpha_F \in \{0.06, 1.0\}$ |
| $\epsilon$ | Parameter mitigating density dependent attenuation of seeds on germination. $\epsilon = 0.2$ |
| $d_F, d_S, d_H$ | Death rates for the flowering plant, seeds/seedlings, & herbivore. $d_F = 0.1, d_S = 0.1, d_H = 0.2$. |
| $h_F, h_2$ | Handling times for herbivory on reproductive adults ($h_F$) and seedling predation ($h_2$). $h_F \in \{0.5, 1.0\}$ & $h_2 \in \{0.5, 1.0\}$ |
| Sub-Function | Definition |
| $\delta$ | Density of seeds produced |
| $\gamma_{12}$ | Density of seeds ($S_1$) maturing to seedling stage ($S_2$) |
| $\gamma_{2F}$ | Density of seedlings ($S_2$) maturing to fecund stage ($F$) |
| $\theta_F$ | Density of adults ($F$) lost to consumption |
| $\theta_2$ | Density of seedlings ($S_2$) lost to consumption |

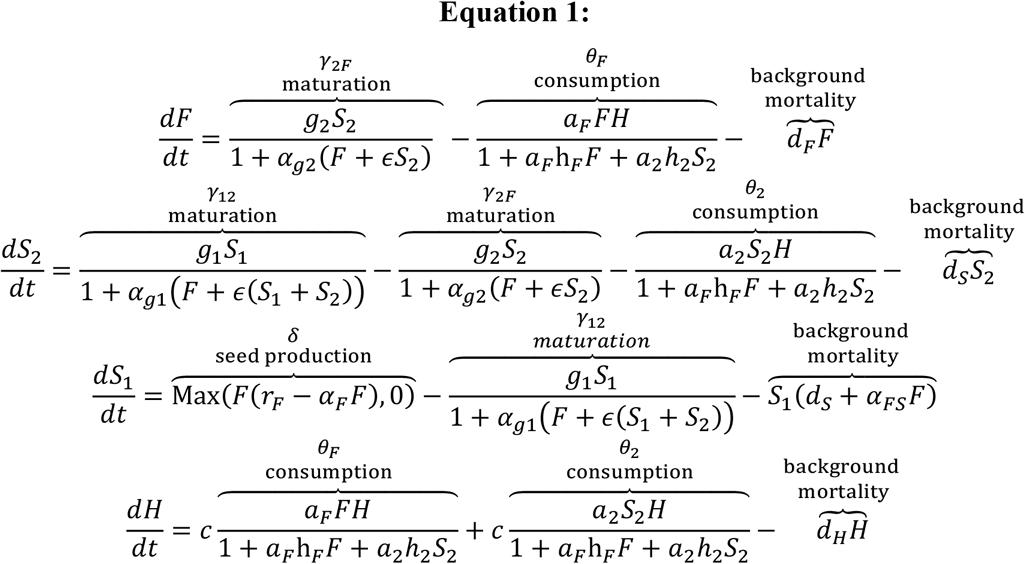

### Simulation Design

Simulations and numerical analyses were done in Mathematica 11. We focus analysis on three specific demographic parameters and two trophic parameters. The demographic parameters are germination rates of seeds into seedlings (*g*_12_), seedling maturation rates into fecund adults (*g*_2*F*_), and seed production rate by fecund adults (*r*_*F*_). The trophic parameters are the attack rates of herbivores (*H*) on seedlings (*a*_2_) and fecund adults (*a*_*F*_). These five rates represent both demographic and trophic flow, a functional basis for studying how plant ontogeny interacts with trophic dynamics (Fig 1a).

We informed the ranges of demographic parameter values simulated in our parameter sweeps using empirical data of three-stage plant population dynamics ^21^. Reproduction from the third stage (*F*) into the first stage (*S*_1_) displayed a large range of values but was highly left skewed, with ∼87% of values <=10. Our exploratory analysis was done with *r*_*F*_<10 and detailed analysis held at *r*_*F*_<=4 (Appendix S1.2; Fig S1a). Stage transitions, on the other hand, are much more evenly distributed between 0 and 1 (Fig S1b&c), so analysis considered 0.12<*g*_12_, *g*_2*F*_ <=0.88. Despite the skewness in the data, analysis reveals no correlation between any demographic rates, indicating no appreciable covariation between parameters (Appendix S1.3; Fig S2). In demographic terms, no covariation between these rates means there was no clear evidence of tradeoffs such that, for example, a higher seed production rate necessitated lower germination rates, or direct correlative relationships between the stage transition rates. The range of herbivore attack rates was chosen heuristically (see Table 1). The lack of any apparent restrictions or relationships with parameter values spurred a broad factorial parameter sweep across all five parameters via cluster computing. We duplicated the five parameter factorial sweep across a range of handling times and degrees of density dependence to test for ubiquity in qualitative results, producing nearly 5.5 million simulations.

### Simulation Analysis

Simulation output measured various factors (Appendix S1.3), but focused on local stability indicators (maximum eigenvalues) signifying stable trajectories (Fig 1b) or persistent oscillations (Fig 1c). Stability indicators are a convenient and fundamental description of the ecological dynamics resultant from each parameter combination. There are multiple machine learning approaches that can provide useful inference to high dimensional nonlinear analysis. We used random forests because they are: 1) relatively easy to implement given their low set of tuning parameters, 2) relatively easy to interpret given their foundation in classification and regression trees, 3) frequently used for feature/predictor selection, 4) flexibly applied across numerous biological fields, 5) and can readily be applied to both categorical and quantitative data ^30,31^. Using the randomForest package in R ^32^, our five simulation model parameters (Fig 1a) functioned as features/predictors with local-stability indicators serving as our predicted variables (Appendix S2). To avoid terminology confusion, we refer to simulation model parameters in general as “parameters” and refer to them specifically as “features” when used as random forest inputs (i.e., independent variables). We then use the term “predictor” only to specifically refer to independent predictors used in our liner models.

For categorical tasks we used a simple indicator, locally stable or unstable. For regression tasks we used the model equilibria’s eigenvalues. We trained random forests using hold out cross validation methods. As a default, random forest parameter “mtry” was set at floor(sqrt(p)) for categorization tasks (stable vs unstable) and floor(p/3) for regression tasks (max eigenvalue) where p=# of features (see Appendix S2.1). Instances where a different p produced better results are noted below. The random forest parameter “ntrees” (No. of trees) was varied from 300-600 with little to no effect on performance. Note these random forest parameters specifically refer to random forest formulation and are not related to the simulation model parameters. We measured random forest performance on validation/test data using area under the receiver operating characteristic curve (AUC; pROC package) for categorization tasks and RMSE for regressions. We measured Variable Importance of individual features in forming predictions with Mean Accuracy Decrease and Mean Increase in MSE for categorization and regression respectively (see Appendix S2; Fig 1). An example of our use of random forests in our analysis is available in SI File 2.

Our five simulation model parameters (3 demographic and 2 trophic rates; see Table 1) served as random forest input features. We determined the degree to which they were independent or interdependent on one another in their effect on random forest predictions with the H-statistic ^33^. We examined the specific details of these interacting features’ effects on trophic dynamics with partial dependence (PD) plots and Individual Conditional Expectation (ICE) curves using the iml package in R ^34^. These results served as the basis for our graphical analysis.

## Results

### Random Forest: Feature Importance & Interactivity

Our random forests produced highly accurate predictions of local stability when trained on model output from the full dataset (e.g., AUC=0.998 across all 5 parameters, see Fig 1d) and all tested subsets. Running random forests on the full results set with all five parameters as predictors indicated both demographic and trophic rates were important to understanding resultant model stability. Moreover, results reveal that whether in multi-stage (red line; Fig 1d) or single stage herbivory (e.g., *a*_2_=0,*a*_*F*_>=0; blue line Fig 1d), parameters’ contribution to predictive power is related to their interactivity with other parameters (blue line; Fig 1d). Note, a similar analysis with *a*_2_>0 & *a*_*F*_=0 is not possible because this type of herbivory is always stable.

This interactivity was apparent in our attempts to understand how our specific parameters affected the behavior of our model in Eq. 1 via studying their effects as features in driving random forest predictions. Initial investigations into individual feature effects revealed that the effect of any single feature (parameter) on trophic dynamics could change substantially based on the values of our other features (parameters). Specifically, the average marginal effects (e.g., PD plots; Fig S3) on simulation dynamics belied a high degree of variability in feature effects throughout the simulation data (e.g., ICE plots; Fig S3).

Breaking down results into further subsets of set specific attack rates with varying demographic rates revealed that this variability in feature effects was largely based on the changes in feature importance and effect over different allocations of herbivory on ontogenetic stages. This breakdown affected the relationship between importance and interactivity (Fig 1d) such that it was inconsistent but still visible in aggregate across our simulation parameters (Fig 1e & 1f). Figures 1g, 1h, and 1i depict how different allocations and intensity of herbivory change the influence of each demographic parameter in driving model stability.

In order to investigate how interactions between demographic rates and stability change as a function of different attack rates, we performed a hierarchical analysis across different allocations of herbivore attack rates. By limiting the number of varying features, we use multivariate analysis to develop a fuller understanding of dynamics in subsections of the data which functioned as a scaffolding for further investigation. In this case, we developed and used an understanding of single-stage herbivory as a basis to study single-stage dominant herbivory (Fig 2), which then leads us to a better overall understanding of our system’s trophic dynamics.

**Figure 2:**
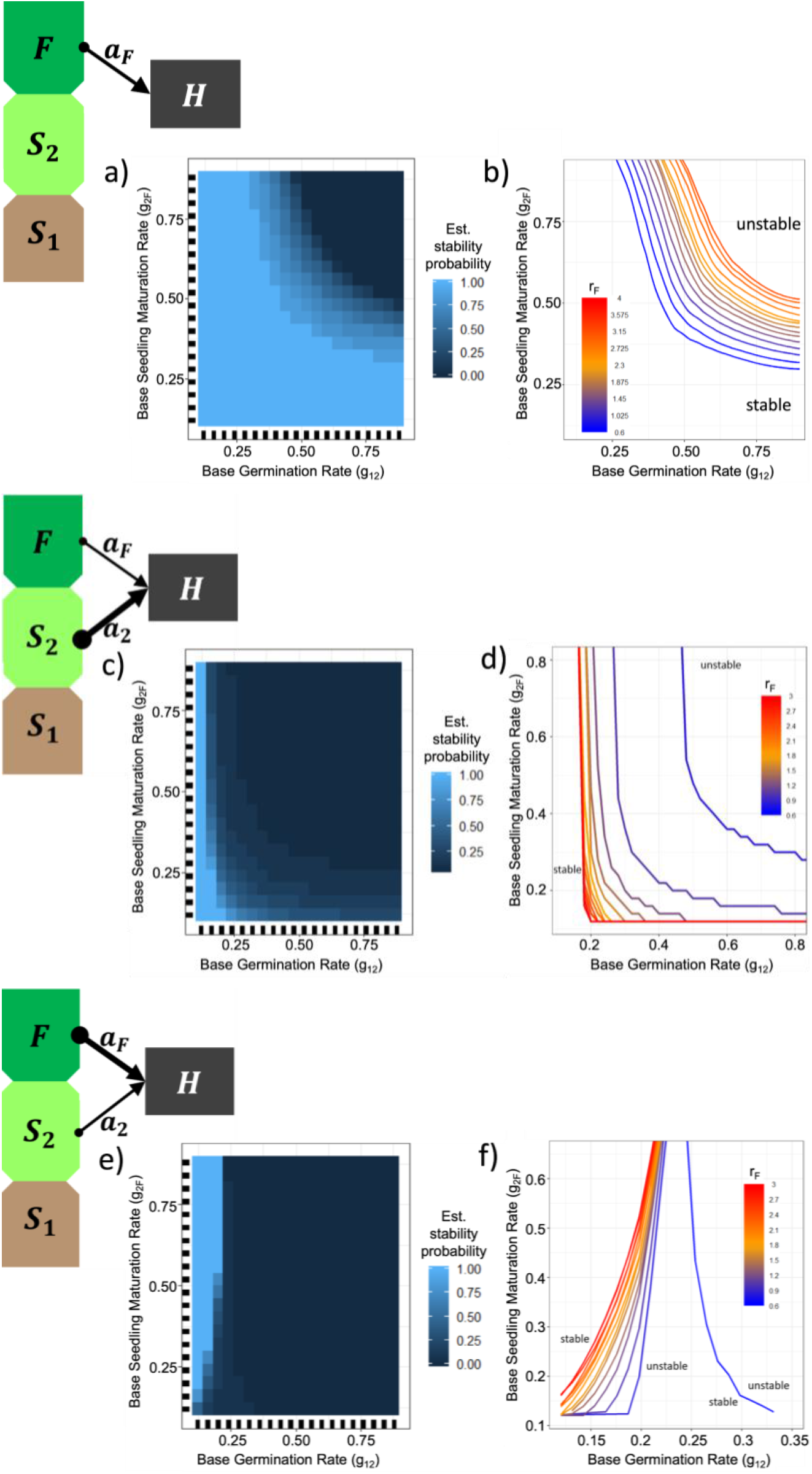
Interactive feature effects on model behavior. Across different herbivory allocations, partial dependence (PD) plots (Fig a,c,e) show interactive effects between maturation rates on categorical simulation stability. Threshold plots (Fig 2 b,d,f) extend this analysis to include gradations of seed production rates. a &b) Herbivory allocation *a*_*F*_=1.0 & *a*_2_=0.0. a) Partial dependence plot shows probability of stability across all values of *r*_*F*_. b) Threshold plot shows the location of the threshold between stable and unstable dynamics in {*g*_12_,*g*_2*F*_} parameter space as a function of seed production levels (*r*_*F*_). c & d) Herbivory allocation *a*_*F*_=0.2 &*a*_2_=1.0. c) Partial dependence plot shows probability of stability across all values of *r*_*F*_. d) Threshold plot shows the location of the threshold between stable and unstable dynamics in {*g*_12_,*g*_2*F*_} parameter space as a function of seed production levels (*r*_*F*_). e & f) Herbivory allocation *a*_*F*_=1.0 & *a*_2_=0.0. e) Partial dependence plot shows probability of stability across all values of *r*_*F*_. f) Threshold plot shows the location of the threshold between stable and unstable dynamics in {*g*_12_,*g*_2*F*_} parameter space as a function of seed production levels (*r*_*F*_).

### Single stage consumption

In the case of the seedling-only herbivore (S2; via *a*_2_>0 & *a*_*F*_=0), all simulations produced stable trophic dynamics. This occurs because density loss in the seedling stage means more juveniles never reach maturity and reproduce themselves (see ref ^18^). This essentially reduces the effective reproduction rate, limits the reproductive plant density, and decreases resources available to the herbivore (similar to lowering intrinsic reproduction in the classic Lotka-Volterra model). In fact, seedling herbivory only induced oscillations at higher handling times, a common effect of high handling time (results not shown).

On the other hand, concentrating consumption on the fecund stage (*F*) can induce both stable and oscillating trajectories (Fig S4). Consumption of *F* does not induce the same regulation of reproductive potential that stabilizes under seedling-only consumption, and so is vulnerable to boom/bust populations cycles. We chose the two most important and interactive parameters (*g*_12_ and *g*_2*F*_; Fig 1e & 1f) in order to search for dominant effects on model behavior and their interactions. This allowed us to create a two-dimensional PD plot ^35^. depicting the estimates of marginal effect of each parameter on random forest predictions, which in this case is categorical stability (Fig 2a). We can see that stability estimates are increased by lowering either or both per-capita germination and/or maturation rates (*g*_12_ and *g*_2*F*_). Demographically, reduced maturation rates shift the ratio of plant population density across its ontogeny, creating a larger juvenile population shielded from consumer pressure. Trophically, this restricts resources for the herbivore, thereby limiting losses in plant density due to herbivory (*θ*_*F*_) relative to the overall plant density.

This mechanism is so influential in determining trophic dynamics, its effect on stability is statistically detectable pre-simulation via equilibrium values. Losses in plant density due to herbivory are labeled under brackets in Eq 1 as *θ*_*F*_ and *θ*_2_, which we can represent as *θ*^∗^_*F*_ and *θ*^∗^_*F*_ at equilibria. Relative to overall plant density we can define a ratio for plants of consumptive losses to total density (L:D ratio) such that:

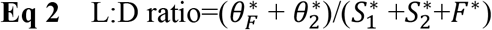

When applied as a predictor variable on the same adult-herbivory subsection presented in Fig 2a via a simple linear regression, we can see that L:D ratio alone explains ∼45% of the variance of the maximum eigenvalue in simple linear models (F-statistic: 4578 on 1 and 5598 DF, p-value: < 2.2e-16) and produces an AUC score of ∼0.83 when predicting categorical stability. Comparatively, our random forest using simulation parameters produced an AUC of 0.98, making it clear that our L:D ratio mechanism explains some but not all the variance in stability outcomes.

Our PD plot (Fig 2a) shows that as both *g*_12_ and *g*_2*F*_ increase, predictions gradually shift from stable to unstable. Based on this observation, we can make a “threshold plot” which depicts thresholds between our categorical variables, stable and unstable behavior, as a function of a third yet unexamined parameter, which in our case is seed production, *r*_*F*_ (Fig 2b). Plotting the thresholds between our stability categories shows a similar dynamic between *g*_12_ and *g*_2*F*_ as seen in the PD plot. It also reveals that the gradual changes seen in the PD plot were in fact a function of the rate of seed production, r;_*F*_, where higher seed production supports stability at higher maturation rates. This is striking given that increased resource production is generally a destabilizing influence in the traditional Lotka-Volterra formulation. Increases in seed production are also related to increases in L:D ratio (Fig S5), so the stabilizing effect of *r*_*F*_ must be coming from a different mechanism. Using a similar analysis on pre-simulation equilibrium values as was done with L:D ratio, we can integrate *δ*^∗^ (see Eq. 1 & description in Table 1) into our regression given its clear connection to rF values. Doing so raises our explanatory power in our regression (R^2^=0.95 in this subset of data) and our predictive power (AUC=0.98), giving us comparable results to our random forest. However, this explains very little of the ecology given that δ^∗^ is largely just the immediate realized effect of *r*_*F*_ and doesn’t describe any other changes in system behavior.

Examining the effects of *r*_*F*_ on the rest of our system shows that *r*_*F*_ is also positively correlated with functions 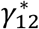 and 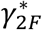 (see definition and description of these functions in Eq. 1 and Table 1, respectively; Fig S5). This implies a stabilizing effect of high densities of maturing seedlings, which seems unintuitive given the stabilizing effect of lowering baseline maturation rates, *g*_12_ and *g*_2*F*_. The key difference is the mechanistic difference between increasing 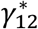 and 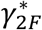 via baseline maturation rates (*g*_12_ & *g*_2*F*_) vs increasing overall seedling density via increases in seed production (*r*_*F*_) (see Appendix S3.2 for more details). Increasing 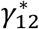 and 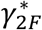via baseline maturation rates (*g*_12_ & *g*_2*F*_) changes the density distribution ratio in favor of *F* (Fig S6a), inducing boom-bust cycles and oscillations as more plant individuals move into the adult stage and are consumed. On the other hand, increased seed production (*r*_*F*_) increases overall plant density, which increases density dependent limitations on maturation, shifts the range of potential density distributions in the plant population, and saturates the younger stages, *S*_1_ and *S*_2_ (Fig S6b). In adult-only herbivory, these saturated stages face no consumer pressure and therefore act as a more immediate reservoir to replace adults lost to consumption by *H*. The functions 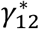and 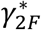 increase because there are simply more seeds and seedlings maturing. Therefore, even though increasing plant density may lead to higher overall consumption, the reservoir of density in younger stages titrates into adult stages due to consumption, raises the trough of oscillations as *r*_*F*_ (and 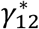 and 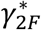) increases (Fig S7) and ultimately leads to a stabilizing consumer dynamic.

This description offers a fuller ecological explanation of how the distribution of density across plant stages mediates herbivore resource availability and drives the observed parameter context-dependence. Specifically, we observe two ways that shifting internal plant demography can promote stability. First, we observe that lowering the average per-capita maturation rates (*g*_12_ and *g*_2*F*_) shields plant density in younger stages. This sequestration of plant density stabilizes by directly restricting resource availability for the herbivore and preventing overconsumption. Second, we observe that raising the intrinsic seed production rate (*r*_*F*_) saturates plant density across the plant population which results in increased resource availability for the herbivore. Despite this increase in resource availability, the system is stabilized by density-dependent limitations on maturation and a robust supply of replacement plant density in younger stages which prevents overconsumption of the adult stage. The observation that increasing resource availability for the herbivore can be both stabilizing (via *r*_*F*_) or destabilizing (via *g*_12_,*g*_2*F*_) depending on the specific parameters underlies much of the parameter context dependence detected by our initial random forest (Fig 1d).

Integrating the 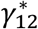 and 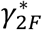factors into our earlier linear regression analysis (with partial least regression due to collinearity between our variables) shows that our three factors (L:D ratio, 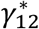 and 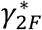) explain 91% of the variance of the maximum eigenvalue (from Fig 2a&2b) and matches our expected direction of effect (shown in Fig 3a). Predictive power increased as well (AUC: 0.98 for categorical stability; RMSE 0.003 for regression on max eigenvalues).

**Figure 3:**
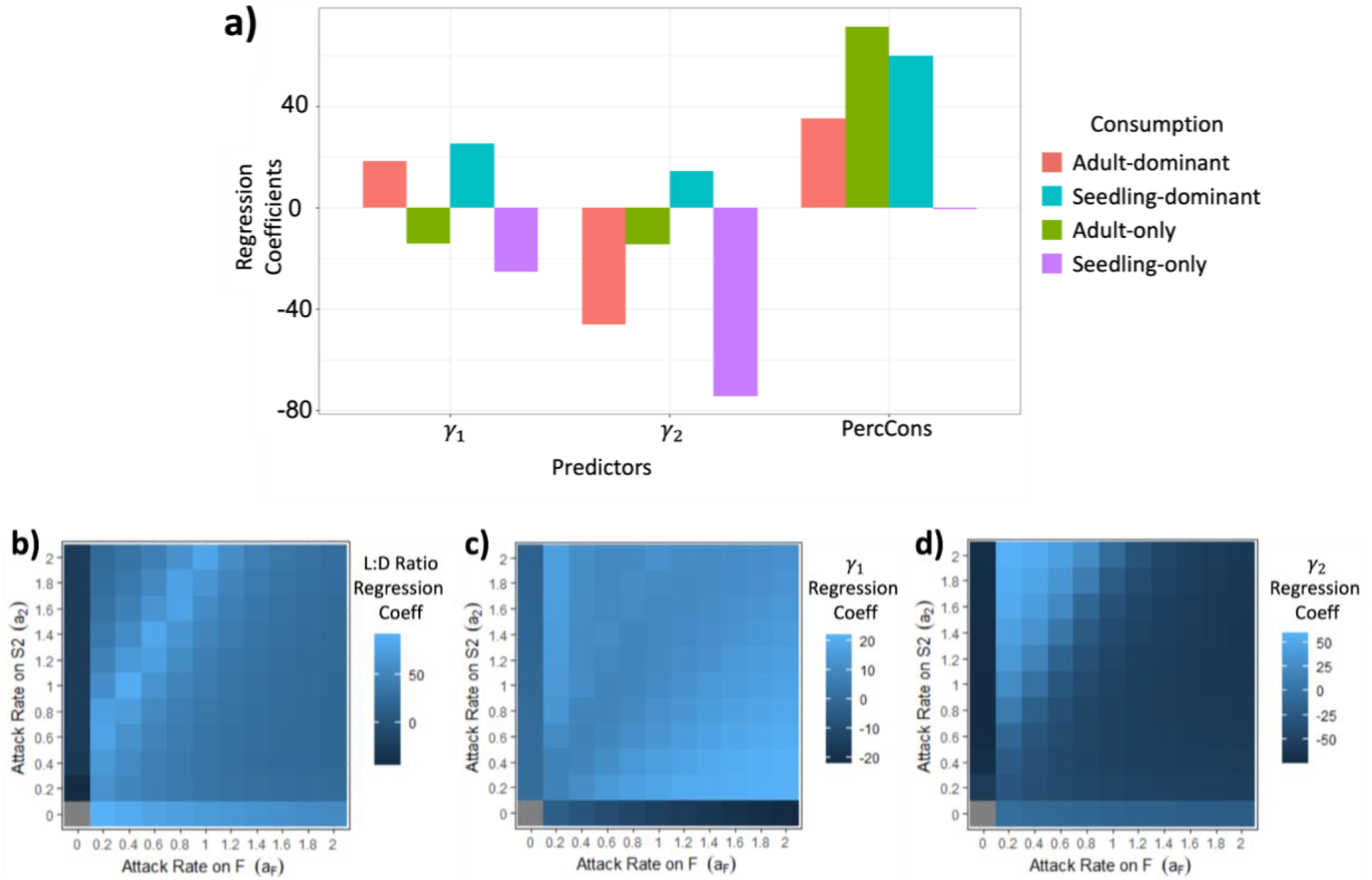
Ecological drivers of stability. Ecological factor effects on stability via coefficients from partial least squares regression of ecological factors versus maximum eigenvalue. Regressions are run for multiple herbivory allocations. a) Bar graph showing specific changes in effect on maximum eigenvalue (regression coefficients) for each ecological factor across the 4 specific herbivory allocations investigated in-depth in the Results. b-d) Range of effects for each ecological factor on stability shown via heatmaps of regressions coefficients on maximum eigenvalue across all specific combinations of herbivory on the adult and seedling stages. Note, the gray square when both attack rates are set to 0 indicates no data given the lack of consumption.

### Multi-stage consumption

Having established baseline dynamics, we move into multi-staged consumption by supplementing single-stage consumption with ancillary consumption on the complementary stage. Specifically, we start by supplementing a seedling-oriented herbivore with limited attack on adults (*a*_2_=1 & *a*_*F*_=0.2) while revealing the importance of interactivity in parameter effects on understanding model behavior. Using this subset of simulation data, we trained a random forest and tested its predictive accuracy. As expected, random forests are sufficiently flexible to maintain high predictive accuracy despite these changes (AUC: 0.98 for categorical stability; RMSE: 0.0001 for regressions of max eigenvalue).

Simulation results reveal that the stability of the seedling-only consumer is vulnerable to destabilization from even limited multi-stage consumption (Fig 2c&2d). PD plots offer some ecological explanation in showing that oscillations can still be stabilized by restrictions in maturation rates (Fig 2c). Extending this analysis with our threshold plots we see that the effect of rF has effectively been reversed (Fig 2d). Overall then, the slight addition of adult consumption institutes a relationship between *g*_12_ and *g*_2*F*_ that mimics an adult-only herbivore but with the caveat that higher *r*_*F*_ values require substantially more restricted maturation to stabilize dynamics.

We can explain this result using our earlier ecological factors (L:D ratio, 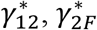). Compared to the adult-only herbivore, we can see that in the seedling-dominant herbivore,raising *r*_*F*_ has much higher proportional effect on L:D ratio compared to 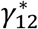 and 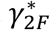 (Fig S8). This is because L:D ratio now consists of consumption on both stages and the seedling stage can no longer act as a saturating reservoir with increased seed production. Additionally, increasing either 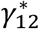 or 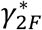 via *r*_*F*_ (higher density) induces higher cumulative attacks (*θ* values; see Eq 1 & Table 1) on both stages. This becomes clear when we regress our ecological factors on our eigenvalue data from this subset of herbivory data. We see that the effects of 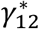 and 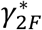 have switched from stabilizing to destabilizing (i.e., raising the max eigenvalue; Fig 3a). Despite these changes, our linear model using our ecological factors still performs comparatively well to our random forest predictions (AUC: 0.99 for categorical stability; RMSE 0.002 (R2=0.97) for regression on max eigenvalues).

We tested this further by supplementing an adult-oriented herbivore with a limited attack on the seedling stage (*a*_2_=0.2, *a*_*F*_=1). Once again, we trained and validated a random forest on this subset of herbivory data and once again the random forest proved adept in providing accurate predictions (AUC: 0.98 for categorical stability; RMSE: 0.0002 for regressions of max eigenvalue). Similar to before, the addition of only slight amounts of multi-stage consumption qualitatively changes the relationships amongst model parameters (input features) and stability. PD plots show that lower *g*_12_ is again stabilizing. However, it also reveals stability at lower *g1*_2_ values is now more dependent on *higher g*_2*F*_ (Fig 2e). Extending the analysis with our threshold plots again indicates that higher seed production values (*r*_*F*_) limit stable {*g*_12_, *g*_2*F*_} parameter space (Fig 2f).

Similar to seedling-oriented herbivory, this multi-stage consumption limits the function of saturating reservoirs in the seedlings and induces more instances of oscillatory dynamics compared to single stage consumption on the fecund stage. However, using our ecological factors, we can investigate the demographic conditions which still promote stability. The stability found at high *g*_2*F*_ and low *g*_12_ can also be described as high 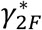 and low 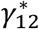 with exact values contingent upon the seed production rate (Fig S9a). These demographic conditions reduce the composition of plants in the seedling stage. This limits herbivore consumption on the seedlings (Fig S9b) and causes the interaction to function more like single stage consumption. This functional similarity to single stage consumption on the adult stage means this promotes the stabilizing effect of a reservoir in the seed bank (low *g*_12_) and a high replacement rate of adults (high *g*_2*F*_). Consequently, simulation results show higher rates of stability at these conditions (Fig S9c) and the coefficients from our partial least squares regression correspond with our analysis (Fig 3a). Additionally, our ecological factors once again not only aid in explaining our results but also perform well as predictors (AUC: 0.99 for categorical stability; RMSE: 0.003 (R^2^=0.99) for regressions of max eigenvalue).

In fact, across all of unique herbivory allocations, our ecological factors perform well as predictors in both categorical (mean AUC: 0.99) and regression based (mean RMSE: 0.002) linear models. This confidence in our predictions allows us to use our partial least squares regression coefficients (on max eigenvalues) to investigate how the effects of each ecological factor change across different allocations of herbivory (Fig 3b-3d). With this we get a detailed view of how plant demography interacts with trophic rates to drive dynamics of our trophic interaction. These results were generally qualitatively consistent across handling times and strength of density dependence (see Table 1) with a notable exception when handling times for seedling consumption are smaller than for adult consumption (see Fig S10).

Finally, expanding back out and considering the full simulation dataset reveals our ecological factors demonstrate predictive accuracy even across all demographic and trophic rates. Using our linear partial least squares model on maximum eigenvalues showed modest success (mean RMSE=0.012 across all permutations of handling time and density dependence). Categorically predicting stability using our factors and the binomial regression was comparatively more successful (mean AUC=0.91 across all permutations of handling time and density dependence).

## Discussion

Increasingly realistic model frameworks in ecology present increasingly complex datasets and novel challenges in analysis. Machine learning algorithms are an obvious candidate for addressing analytical challenges. While machine learning algorithms still have limits in their interpretability, their ability to produce highly accurate predictions bolster their increasing prevalence in ecology ^10^. Stopping at predicting ecological model outcome alone, however, places high credence in simulation model formulation if no mechanistic explanation behind the predicted outcome is possible. Famously, “all models are wrong” ^36^ but their utility lies in their ability to aid researcher’s intuition of complex systems. Therefore, mechanistic explanations of model results are critical to achieving models’ maximum utility in ecology. Accordingly, there is fundamental value in expanding the interpretability of machine learning (e.g., random forests) in studying simulation models which we argue connects to the core utility of modeling in science.

The variety of ecological behavior across organismal ontogeny is an important frontier in ecological modeling ^15^. The context-dependent complexities resulting from even our simple inclusions of ontogeny make clear the need to develop broadly applicable methodologies which improve generalizable ecological understanding. Our random forest output produced clear descriptions of each simulation model parameters’ contribution to predicting simulation behavior and our H-statistic analysis showed that these contributions were largely tied to interactions with other model parameters (both demographic and trophic; see Fig 1). Individual Conditional Expectation plots revealed the extent of this context-dependency (e.g., ICE plots Fig S3), but our goal was to go beyond the well-known ecological axiom that parameter effects are context-dependent. We therefore subdivided our data into its component parts and focusing on the consistently important model parameters and it became clear that different parameters drove model behavior across different herbivory allocations (Fig 1g-1i). By investigating feature interactions using two-dimensional PD plots and then extending categorical analysis with our threshold plots (Fig 2), we were able to move beyond connecting model behavior to parameter values to describing model output via tangible ecological processes.

For ease of communication, we called our ecological processes (L:D ratio, 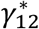, and 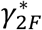) “factors.” Using these factors, we detailed how demographic and trophic rates interact to drive ontogenetically mediated consumer-resource dynamics via a consistent set of ecological mechanisms. These factors were also able to predict model output with comparable accuracy to that of the random forest. Furthermore, despite being found from categorically tasked random forests (stable or unstable), these same factors performed well on continuous predictions (maximum eigenvalues) given the relatedness of each variable.

In expanding upon our work to higher dimensionalities found in food web/network models, several considerations come to mind. First, random forests facilitate intuitive feature selection which allows researchers to deal with higher parameter counts by focusing on the most important simulation model parameters (via accuracy decreases, see Methods). Additionally, subsetting simulation data (as we did above by focusing on specific herbivory allocations) simplifies analysis and further facilitates finding mechanistic drivers of model dynamics. In cases where parameterization is systematically controlled and not varied (e.g., the Allometric Trophic Network), analysis of model behavior can easily be refocused onto quantitative network properties via random forest inputs.

Finally, while our analysis used equilibria and linear stability metrics, research at the level of food web analysis typically focuses on other metrics of model behavior. This is partially because finding and expressing equilibria in high dimensional models can be difficult. However, it has been shown to be tractable ^20^ when using root finding algorithms which do not rely on explicit parametric expressions ^37^, but instead provide the numerical values of equilibria which would be sufficient for our purposes. On the other hand, other metrics frequently used at the food web/network scale, such as species extinctions ^6^, temporal variation ^38^, biomass production ^13^ can readily fit the random forest framework. Additionally, by establishing threshold cutoffs of interest akin to our dual use of categorical stability and maximum eigenvalues, researchers can couple categorical and regression tasks to maximize their ability to discover mechanistic drivers of model behavior via the processes detailed here.

We do not claim our exact methodology will be entirely applicable for all models or questions, but instead aimed to present an example process extending the analytical power of random forests of prediction to mechanistic explanation. In the case of stage-structured ontogeny, our process revealed that complex interactions amongst parameters could be consolidated into key ecological mechanisms. These mechanisms explained how trophic interactions mediated by plant ontogeny can produce disparate results compared to traditional model structure. The extent of these differences exposes the need to further integrate unique sources of ecological complexity to better understand drivers of community dynamics.

## Acknowledgements

This research was supported by National Science Foundation grants DEB-1834497 and DEB-2129757 to F.S.V. S.S. was partially supported by the UC Davis Grant for Advancing Sustainable Development Goals, Global Affairs as awarded to F.S.V and the Sustainable Oceans National Research Trainee Fellowship (NSF award # 1734999) as awarded to S.S. This research was supported in part through computational resources and services provided by Advanced Research Computing (ARC), a division of Information and Technology Services (ITS) at the University of Michigan, Ann Arbor.

## Author Contributions

S.S., P.G., F.S.V. conceived and conceptualized this project. S.S. & P.G. developed the simulation model. P.G. conceived and implemented the random forest analysis and extension. P.G. &S.S. developed and implemented the linear analysis. P.G. &S.S. wrote the first draft of the manuscript. SS. PG. &FSV contributed to the writing of the following versions.

## Data availability

All COMPADRE data used in our model formulation is available on the COMPADRE platform and the exact specifications are reproducible using Supplementary File 1: CompadreAnalysisRMD.html. Simulation data used in our analysis is available as a zipped file online here.

## Code availability

The simulation model (Eq 1) as well as R scripts for all random forest analysis, main text figures/analysis, and supplementary figures will be made available at https://github.com/prglaum/PlantOntogenyDynamics upon completion of peer review and publication processes. Random forest analysis is available as an R Markdown file in Supplementary File 2 (randomForestCode.html) and as an interactive app for readers at https://prglaum.shinyapps.io/simData-randomForestRMD/.

