## Appendix for "Through the Random Forest: Ontogeny as a study system to connect prediction to explanation"

### S1 - Model development

In this section we use the online database COMPADRE to investigate model structure, plausible parameter values, and simulation design. R scripts to run our analysis are presented in SI github. The analysis presented below was completed in the later part of 2021 and COMPADRE is a growing database. Therefore, attempts to recreate our analysis at a later date are likely to reflect changes made to the database.

#### S1.1 - Model structure

Three distinct stages were used as the ontogenetic structure of the plant population in our model: fecund adults ($F$), non-fecund seedlings ($S_2$), and the seed bank ($S_1$) (Fig 1). While no single ontogenetic structure can perfectly represent all plant species, three stages is a suitable baseline for the purposes of this study. First, a three-stage structure creates a tractable dimensionality in our model which allows for an in depth analysis of interdependent dynamic effects born out of interacting demographic/ecological rates. Importantly, this allows us to more clearly graphically represent the consequences and details of interacting rates which would be less apparent in higher dimensional ontogenetic formulations. Second, using the growing global database of stage-structured plant demographic data, COMPADRE[39], we can see that the three stage ontogeny is well-represented in plant taxa empirically. Of all the documented plant taxa within the COMPADRE database, three-stage plant structures represent roughly 34% of families, 18% of genera, and 14% of species. This includes abundant, species-rich, economically important, and geographically wide-spread plant families such as *Asteraceae*, *Brassicaceae*, *Orchidaceae*, *Rosaceae*, etc.

The plant population in the model experiences density dependent restrictions on the production of seeds via the maximum function which restricts seed production to a minimum of zero in relation to the density of $F$. Stage transitions are also modified by density dependent pressure with limited density dependent effects from "younger" via the parameter $\epsilon$ (see Eq1 & Table 1). Finally, consumer pressure from the herbivore population ($H$) can differentially focus its herbivory on either the seedling stage ($a_2 > 0$, $a_F = 0$), the fecund adult stage ($a_2 = 0$, $a_F > 0$), or both ($a_2 > 0$, $a_F > 0$). Herbivory occurs under a Type II functional response on each stage, where consumption on each stage is affected by the handling time required to consume both stages. All stages experience a background mortality rate. Seeds ($S_1$) experience a low level amplification of background mortality linked to adult ($F$) density under the assumption that sufficiently high populations of conspecific mature plants reduce resources for seeds (through lack of nutrients, shading, etc.)[40] or increase frequencies of exploitative interactions not explicitly unaccounted for in the model (e.g. soil pathogens)[41,42]. Parameters mediating all density dependent effects and trophic interactions are listed and described in Table 1.

$$\frac{dF}{dt} = \overbrace{\frac{g_2 S_2}{1+\alpha_{g2}(F+\epsilon S_2)}}^{\substack{\gamma_{2F}\\ \text{maturation}}} - \overbrace{\frac{a_F F H}{1+a_F \mathrm{h}_F F + a_2 h_2 S_2}}^{\substack{\theta_F\\ \text{consumption}}} - \overbrace{d_F F}^{\substack{\text{background}\\ \text{mortality}}}$$

$$\frac{dS_2}{dt} = \overbrace{\frac{g_1 S_1}{1+\alpha_{g1}\big(F+\epsilon(S_1+S_2)\big)}}^{\substack{\gamma_{12}\\ \text{maturation}}} - \overbrace{\frac{g_2 S_2}{1+\alpha_{g2}(F+\epsilon S_2)}}^{\substack{\gamma_{2F}\\ \text{maturation}}} - \overbrace{\frac{a_2 S_2 H}{1+a_F \mathrm{h}_F F + a_2 h_2 S_2}}^{\substack{\theta_2\\ \text{consumption}}} - \overbrace{d_S S_2}^{\substack{\text{background}\\ \text{mortality}}}$$

$$\frac{dS_1}{dt} = \overbrace{\mathrm{Max}(F(r_F - \alpha_F F), 0)}^{\substack{\delta\\ \text{seed production}}} - \overbrace{\frac{g_1 S_1}{1+\alpha_{g1}\big(F+\epsilon(S_1+S_2)\big)}}^{\substack{\gamma_{12}\\ maturation}} - \overbrace{S_1(d_S + \alpha_{FS} F)}^{\substack{\text{background}\\ \text{mortality}}}$$

$$\frac{dH}{dt} = c\overbrace{\frac{a_F F H}{1+a_F \mathrm{h}_F F + a_2 h_2 S_2}}^{\substack{\theta_F\\ \text{consumption}}} + c\overbrace{\frac{a_2 S_2 H}{1+a_F \mathrm{h}_F F + a_2 h_2 S_2}}^{\substack{\theta_2\\ \text{consumption}}} - \overbrace{d_H H}^{\substack{\text{background}\\ \text{mortality}}}$$

S1.2 - Model Parameters

The COMPADRE database also allows for further insight into our model. We can use the available data to inform certain model parameters. The COMPADRE reproductive and stage transition rates mirror our demographic parameters. Production of stage 1 individuals from stage 3 corresponds to seed production ($r_F$). Transition from stage 1 to 2 corresponds to seed germination ($g_{12}$) and transition from stage 2 to 3 corresponds to seedling maturation ($g_{2F}$). However, these rates represent data from a range of different studies using a range of experimental, natural, and agricultural plant demographic data. Additionally, these studies involve transition rates which emerge from both competition and trophic interactions. Therefore, measurable field rates are frequently emergent from environmental conditions and are not necessarily the inherent demographic rates. As such, we do not intend to use these empirical rates to determine exact values of our model's demographic parameters. Instead, we use the ranges of each empirical rate to inform plausible ranges for their corresponding model parameter to be analyzed in parameter sweeps of model simulations.

In doing so, we limited our survey of values from each parameter to those greater than 0 as the effect of setting any demographic parameter to 0 in the model is obviously a steady fall to extinction (Fig S1). Starting with rates of reproduction (Fig S1a), we see a large range of reproduction into the first stage from the third, from 0 to roughly 100. However, we limited our preliminary analysis to $r_F < 10$ as 87% of sampled values fall within this range (Fig S1a, Insert). The rates of transition between stages are much more evenly distributed from values >0 to 1. While not a completely even distribution across the range, we take this as sufficient reason to study the full range of values, $0.1 < g_{12} < 0.9$ and $0.1 < g_{2F} < 0.9$. With these results in mind (Fig S1), ranges for these three parameters are provided in Table 1 along with both parameter definitions and the ranges/values used for every model parameter.

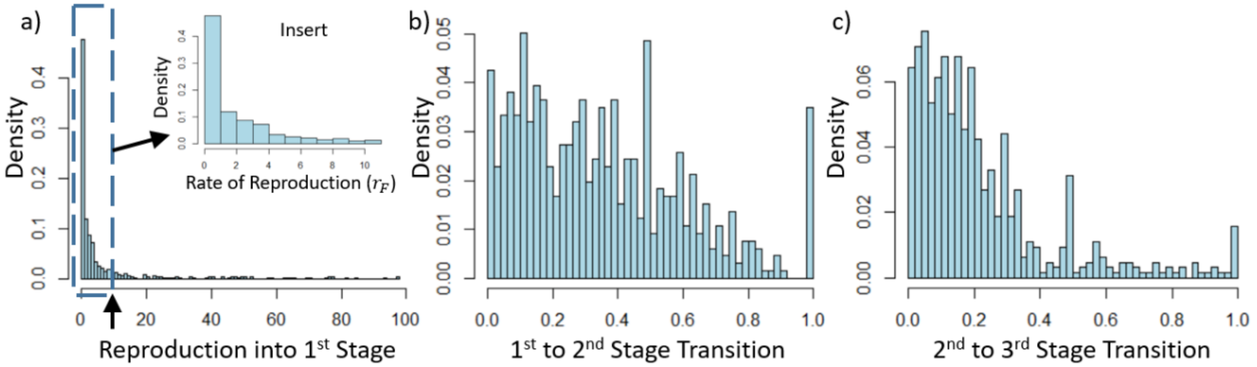


**Fig S1:** Distribution of empirically measured reproduction and stage transition rates from three-stage plant matrix models in the COMPADRE database (2021). a) Reproduction into 1st stage by 3rd stage. b) Transition rates from stage 1 to 2. Transition rates from stage 2 to 3.

S1.3 - Simulation Design & Output

Potentially the most important insight garnered from available data in COMPADRE, we see absolutely no correlation between any of the measured rates. There is no discernable relationship between empirical rates of reproduction and maturation from either stage 1 to 2 or 2 to 3 (visualized in Fig S2a & b). Also, we see no correlation between the stage transition rates (visualized in Fig S2c). The lack of any relationships in the values of any rate prompted a full factorial investigation of the model's demographic rates ($r_F$, $g_{12}$, $g_{2F}$; see Table 1 for value range), without any necessary covariation in parameter values. In other words, all demographic parameter values were factorially tested against each other in a large parameter sweep without the need to change any one parameter value in concert with another. Past work also indicates interactivity between demographic rates and trophic interactions in driving model dynamics[18], so we considered trophic interactions in our simulation design by including rates of herbivory ($a_F$ & $a_2$) into what becomes a five dimensional fully factorial parameter sweep. Herbivore attack rate ranges were chosen heuristically (Table 1). Herbivore attack rates on either consumed plant stage (see Fig 1d-1i) vary factorially in the parameter sweep as herbivores can range from focusing their attack on either stage to splitting their consumption between stages to varying degrees.

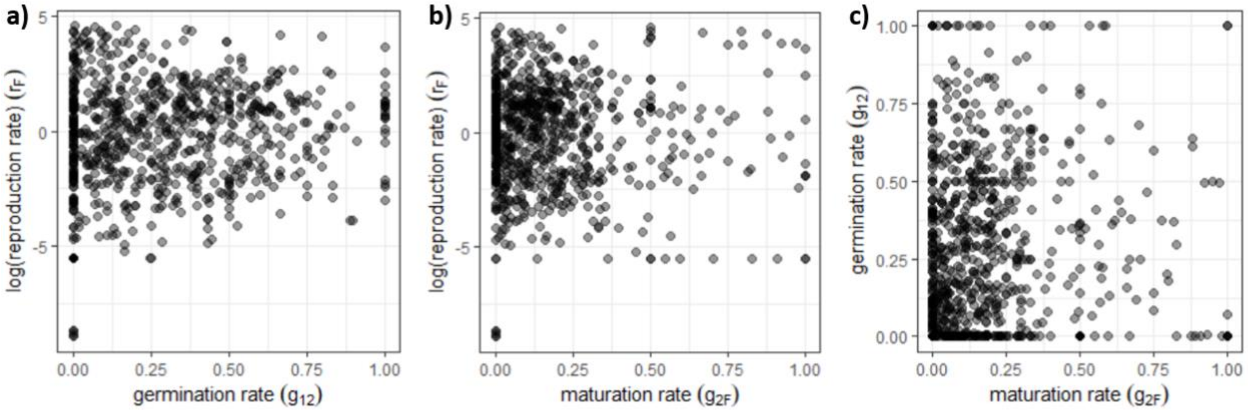


**Figure S2:** Scatter plots showing the lack of correlative relationships between COMPADRE (2021) three-stage reproduction and transition rates. a) Reproduction rates vs transition rates between stages 1 & 2. b) Reproduction rates vs transition rates between stages 2 & 3. c) Reproduction rates between stages 1 & 2 vs transition rates between stages 2 & 3.

As a mode of sensitivity testing, we implemented the five parameter factorial sweep across multiple values for both density dependent parameters and handling times. Density dependent parameters were varied because density dependence mediates stage transitions and therefore ontogenetic dynamics[18]. Handling time was chosen for its role in mediating the trophic connections interacting with plant ontogeny. This created eight unique instances of the full five parameter factorial sweeps (see values in Table 1), producing over 5.5 million unique simulations for analysis.

Each simulation outputs a number of initial conditions and post simulation factors detailing the results of the simulation. Initial conditions include all relevant parameter values and initial time dependent variable densities (i.e., initial population densities such as $F(t)$ at time 0). Post simulation factors include equilibria values, equilibria linear stability, eigenvalues, periodicity, oscillating populations' peak and trough densities, and finally the effective handling time of the herbivore population. The effective handling time of the herbivore population is measured as the denominator of the consumptive interaction in the model.

### S2 - Analyzing Simulation Results w/ Random Forests

Due to the large simulation dataset, we first produced an initial guided analysis using the Random Forest based machine learning algorithm which can achieve high predictive power[43] and has shown success in using permutation techniques to determine how much specific predictors contribute to that predictive ability (e.g., mean accuracy decrease[44]). In our case, our five main parameters (see Table 1) serve as our main random forest features/predictors. Random Forests can predict either categorical or continuous variables and any post simulation factor can function as the predicted variable in the random forest analysis. Our interest in trophic/demographic dynamics led us to use the simulation model's linear stability as our predicted variables. We used a simple indicator, stable or unstable, for categorization random forest tasks and model equilibria's eigenvalues for regression random forest tasks.

S2.1 - Preparing data and training Random Forest

Simulation data was split into “training” and “validation” (sometimes called test) data subsets for hold out cross validation. We first created or “trained” the random forest on the training data. In Random Forests, this process produces a series of unique categorization and regression trees that “vote” on the outcome (our simulation model’s stability) based on the values of any particular inputs (our model parameters). As a default during training, random forest parameter “mtry” was set at floor(sqrt(p)) for categorization tasks (stable vs unstable) and floor(p/3) for regression tasks (max eigenvalue) where p=# of features[45]. Instances where a different p produced better results are noted in the text. The parameter “ntrees” (No. of trees) was varied from 300-600 with little to no effect on performance.

Once we have a trained random forest, we check its performance on the training data via the “out of the box” (OOTB) error rate. Given a sufficiently low error rate, we can begin to investigate feature importance. We measured the importance of individual features/parameters in our random forest with Mean Accuracy Decrease, which measures the loss in predictive accuracy by excluding each feature. The more the accuracy suffers, the more important the variable is for the successful classification/prediction. For a more detailed description of the mechanisms behind creating and training random forests, please see ref 13.

S2.2 - Validating/Testing Trained Random Forest

Sufficiently high performing trained random forests were then used to predict simulation output in our validation data subsets which our random forests had not yet been exposed to. We analyzed the predictive power of categorization tasks by comparing their predicted output with data via the Area Under Curve the Receiver Operating Characteristic curve (AUC) metric (pROC package). In our case, the AUC metric measures how well the models are able to distinguish between stable and unstable results in the validation data subset. It varies between 0 and 1 with 1 indicating better predictions. Regression tasks were judged for accuracy using RMSE on maximum eigenvalue measurements between the predicted eigenvalue output and the simulation data. All AUC and RMSE values are from validation data unless otherwise specified.

S2.3 - Interpreting Feature Effects

Random Forest results can also be further interpreted with the H statistic which functions as a measure of interactivity between features/predictors in driving prediction results[46]. This can be done for each individual predictor as a measure of general interactivity with other predictors or be done with a focus on particular predictors to study direct interactivity between specific predictor combinations. Regardless, higher numbers indicate higher interaction strengths while lower numbers indicate less interaction between predictors. While some have argued that the basic random forest measurements of variable importance (e.g., mean accuracy decrease) “capture” the outcome of interactions between features in predictions, these metrics are not designed to detect interactions per se[47]. Therefore, our use of the H statistic helps us hone in on key interactions, a particularly useful outcome for any researcher looking to implement our methods on a model with a larger parameter set.

While the H statistic provides inference regarding which features have strong interactive effects, other methods are required to determine how features (interacting or otherwise) actually affect the predicted outcomes. To do this we used two analytical techniques in the iml package in R[48]. First, we used Partial Dependence plots (PD plots) to visualize the marginal effect of one or two features on the predicted outcome of our random forests[49]. These PD plots can show whether the relationship between the target and a feature is linear, monotonic or more complex. By focusing on two-feature partial dependence plots, we can also see how these features interact in changing model predictions (e.g., Fig 2). Second, we also used Individual Conditional Expectation (ICE) curves to uncover heterogeneous relationships by showing individual instances of changing a feature’s value at different permutations of the other features[50]. In using these ICE plots, we found quick evidence of the context dependent relationship of features and their effect on model predictions (Fig S3), again helping us determine where interactions matter.

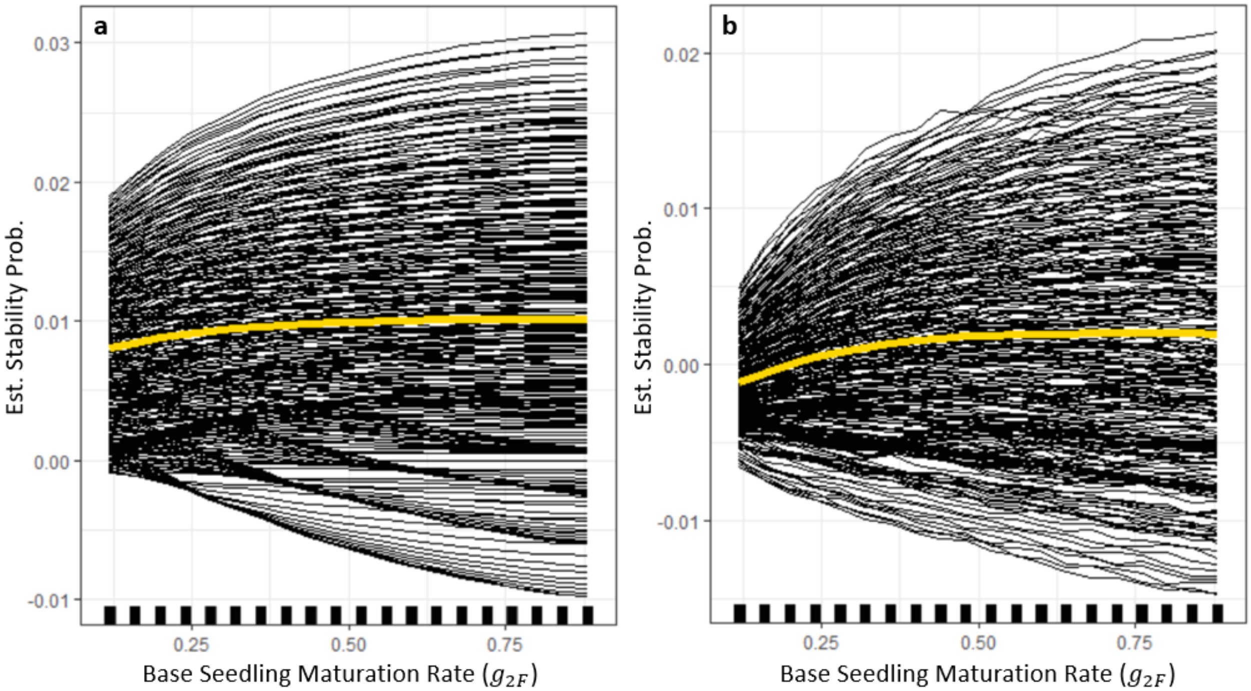


**Figure S3:** a) Example of the high degree of heterogeneity in feature effects on random forest outcome even for set attack rates ($a_2 = 0.2$ & $a_F = 1.0$) using the g2F parameter.
b) Depiction of the high degree of heterogeneity in feature effects on random forest predictions using the g2F parameter as an example.
The yellow line represents the average partial dependence effect (PD plot). The black lines show an Individual Conditional Expectation (ICE) plot; instances of changing g2 in the context of differing subsets of other parameters. Both the quantitative and qualitative range of differences seen in the ICE plot indicate the fidelity lost in only examining the average effects and the need for a more fine scale look at the heterogeneity in effect per parameter. Note, the right ICE plot only shows results for $a_2, a_F <= 1$ in an effort to reduce the number of plotted lines for the sake of visibility.

S2.4 - Additional

Our random forests can produce a highly accurate level of predictive power. These levels of predictive ability can induce questions of data leakage in producing and testing our models. We claim this is not the case here and detail our reasons below. First, the two easiest sources of data leakage are including the target variable as a feature in creating and training our models while the other is accidental inclusion of test/validation data in our training data during model training. Neither of these occurred due to simple due diligence in model creation. Our random forest code can be used to verify these claims. Second, we have no “give away” features which are effectively tied to our target variables. Third, results are not driven by particular outliers and are consistent across different subsets used as training and test/validation datasets. Finally, models’ variable importance changes in explanatory ways across different subsets of data (e.g., different consumption allocations) and our results are supported by our graphical analysis (Fig 2) which does not fall victim to data leakage issues. Therefore, we can have high confidence that our random forest results do not reflect data leakage.

Despite the high levels of predictive power (e.g., results shown in Fig. 1), our random forests did have limits on their immediate interpretability as noted by others (e.g. ref 51). Therefore, we use our random forest models not as end points on their own, but as tools to direct our analysis across such a large amount of simulation data. By determining feature importance, effect, and interactivity, we were able to hone in specific subsections of the simulation model's parameter space and utilize graphical analysis to expand our ecological understanding of our results.

### S3 - Supplementary Analysis Results

S3.1 - Figures

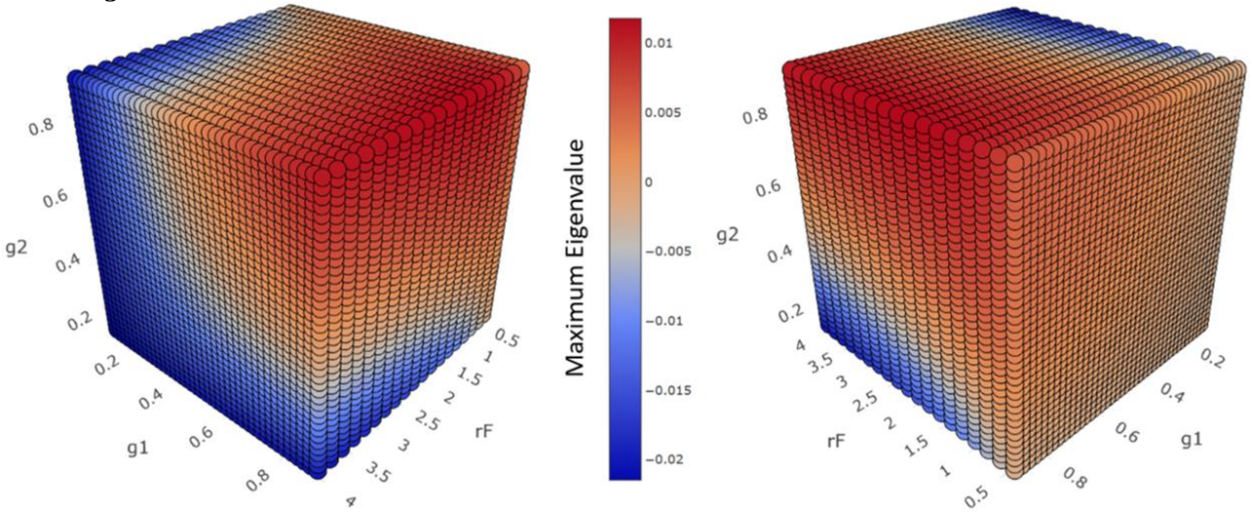


**Figure S4:** Maximum eigenvalue of the plant-herbivore system where the herbivore only eats the adult plant stage ($a_F > 0, a_2 = 0$) across $\{g_{12}, g_{2F}, r_F\}$ parameter space. Maximum eigenvalue here dictates the dynamic stability of the population trajectories of both species in the interaction. Positive values indicate instability and persistent oscillations while negative values indicate damped oscillations to stable population trajectories.

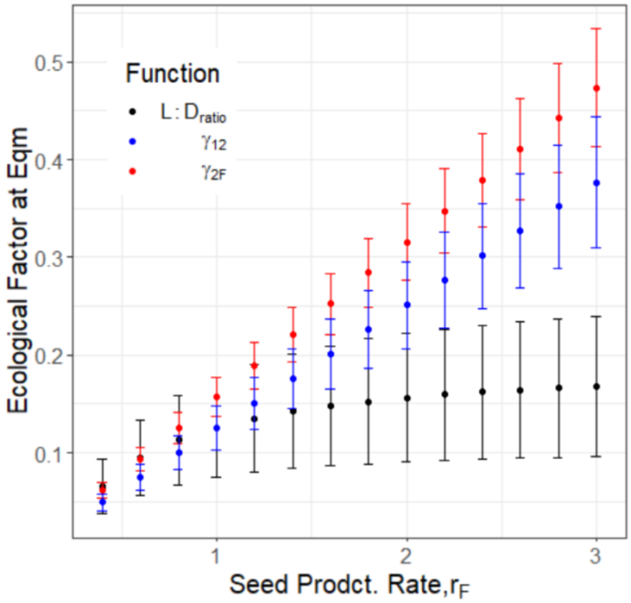


**Figure S5:** Seed production effect on ecological factors with adult-only herbivores. The mean effect of raising seed production broken down to average constituent effects on ecological factors across all other simulation model parameters. Dots represent mean and error bars show standard deviation.

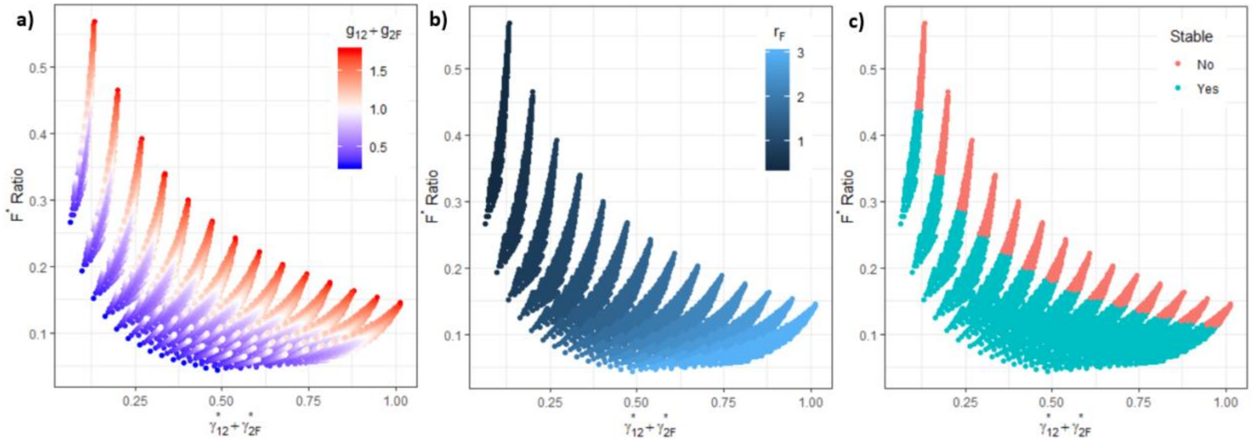


**Figure S6:** Changes in $F^*$ ratio in relation to ecological factors $\gamma^*_{12}$ and $\gamma^*_{2F}$. The y axis, $F^*$ Ratio is the percent makeup of the adult plant stage defined as $F^*$ Ratio = $\frac{F^*}{S_1^*+S_2^*+F^*}$. Color contrast shows constituent changes in a) simulation model parameters $g_{12} + g_{2F}$, b) simulation model parameter $r_F$, c) stability of simulation model.

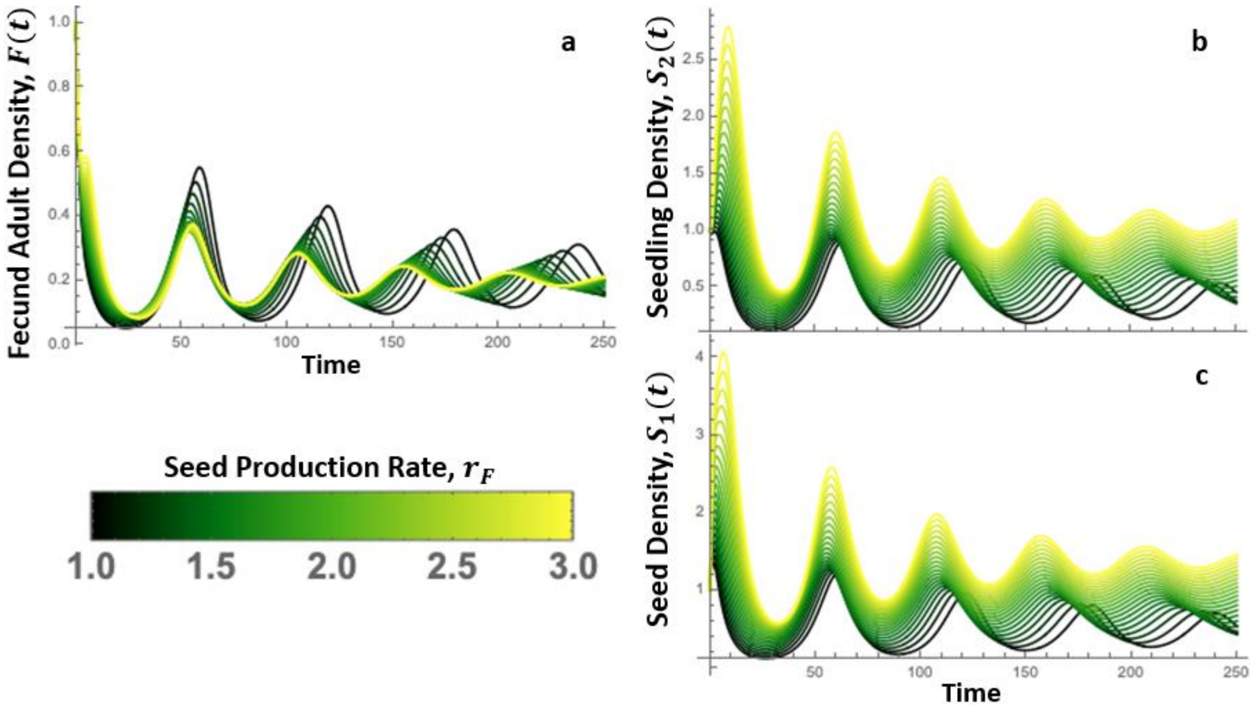


**Figure S7:** Time series of Eq1 with an adult-only herbivore. Model dynamics are shown across gradients of values for seed production ($r_F$) with line color corresponding to the value of the parameters shown in the color legend. Figure displays how raising seed production lifts the minimum density in the oscillating plant population, dampening the oscillations and stabilizing the trophic interaction. Panels display time series for a) Fecund adults, $F$, b) Seedling, $S_2$, and c) Seed bank, $S_1$. Other parameters are as follows: $g_{12} = 0.34$, $g_{2F} = 0.34$, $a_2 = 0$, $a_F = 1$, $r_F = [1,3]$.

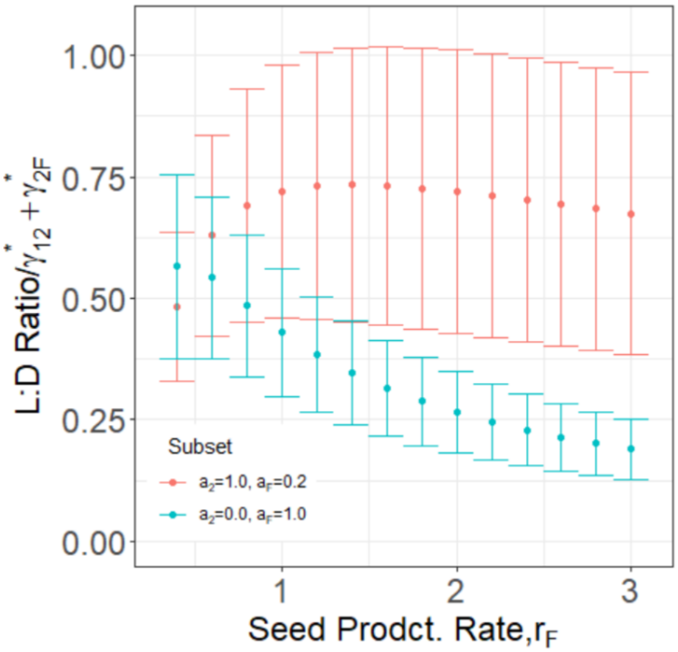


**Figure S8:** The effect of seed production ($r_F$) on our ecological factors. Effects are expressed through the relationship, $\frac{L{:}D\ Ratio}{\gamma_{12}^{*}\ \&\ \gamma_{2F}^{*}}$ to limit clutter of expressing each factor separately. Figure depicts how increasing seed production has a greater positive effect on consumption and consequently $L{:}D\ Ratio$ when $a_2 = 1.0$ & $a_F = 0.2$ than when $a_2 = 0.0$ & $a_F = 1.0$.

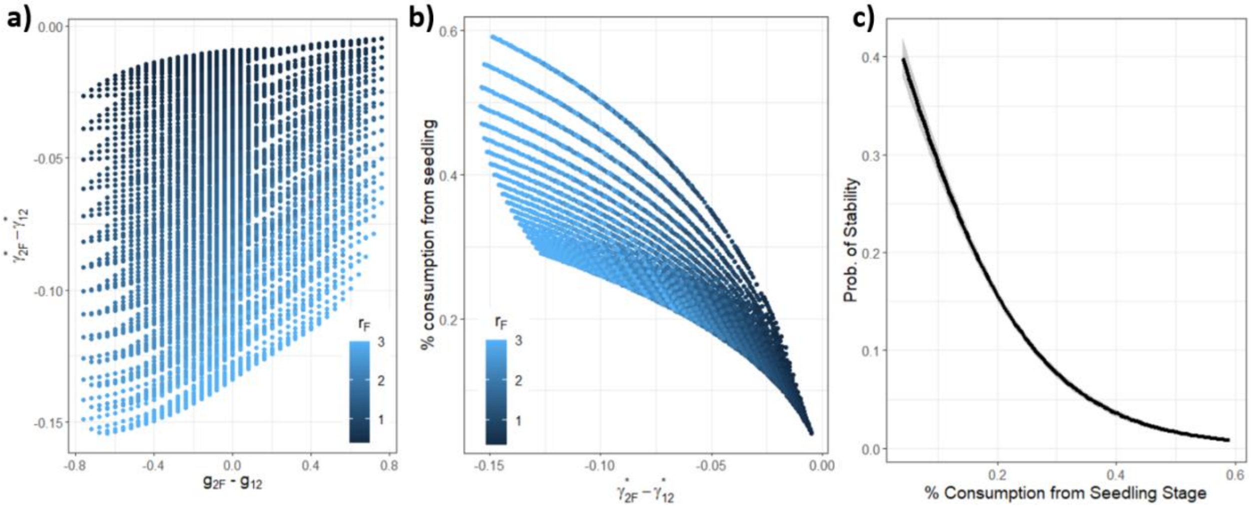


**Figure S9:** Drivers of stabilization in $a_2 = 0.2$ & $a_F = 1.0$ herbivory allocation when $g_{2F}$ is high and $g_{12}$ is low. a) As $g_{12}$ decreases and $g_{2F}$ increases (measured by $g_{2F} - g_{12}$) this consequentially increases seedling maturation while limiting seed germination ($\gamma_{2F} - \gamma_{12}$), reducing the plant density held in the seedling stages. b) As the plant density in the seedling stage decreases, so does the percent consumption of seedlings, measured here by a decrease in $\frac{\theta_2}{\theta_F + \theta_2}$. c) Reducing the percent of seedlings in the diet of the herbivore makes the trophic interaction act more like single stage consumption. This increases the probability of dampened oscillations to stability as shown via a generalized linear model (p<<2e-16, beta=-8.003, Residual deviance: 3815.3 on 5598 degrees of freedom).

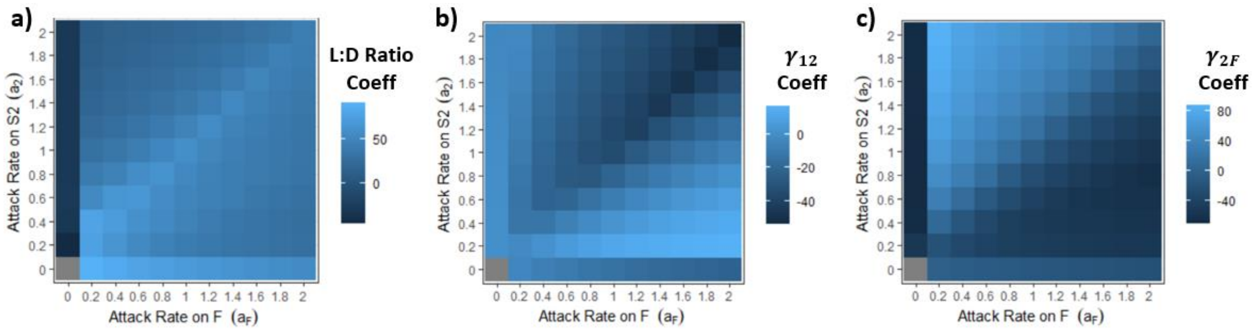


**Figure S10:** Heatmap colors represent ecological effects on stability via coefficients from partial least squares regression of ecological factors versus maximum eigenvalue across all specific combinations of herbivory on the adult and seedling stages when $h_2 = 0.5$, $h_F = 1$ and $\alpha_F, \alpha_{g1}, \alpha_{g2} = 0.6$. The ecological factors are a) L:D Ratio, b) $\gamma_{12}$, & c) $\gamma_{2F}$. Note, the gray square when both attack rates are set to 0 indicates no data given the lack of consumption. The figure depicts how a lower handling time for herbivory on seedlings versus adults can change the dynamic effect of $\gamma_{12}$ such that increased germination can dampen oscillations more consistently across different herbivory allocations.

S3.2 - Description of $g_{12}$ & $g_{2F}$ vs $\gamma_{12}$ & $\gamma_{2F}$:

Although the $g_{12}$ & $g_{2F}$ parameters and $\gamma_{12}$ & $\gamma_{2F}$ ecological sub-functions we labeled "factors" (Table 1) are related, their relationship is not 1:1. Specifically, $g_{12}$ and $g_{2F}$ are parameters in the model representing the per-capita germination/maturation rates of seeds/seedlings. The fixed values of these parameters are assigned for each simulation and are not dynamic. These parameters represent the rate of maturation without density dependent effects.

The $\gamma_{12}$ and $\gamma_{2F}$ sub-functions/components in the model represent the density of seeds/seedlings that germinate/mature to the seedling/adult stages over a timestep. The values of these sub-functions are an emergent property of the model resulting from the interaction between herbivore consumption and internal plant demography. For example, the parameter $r_F$ affects the output of $\gamma_{12}$ and $\gamma_{2F}$. Unlike $g_{12}$ and $g_{2F}$, (which are fixed parameters), $\gamma_{12}$ and $\gamma_{2F}$ may increase with $r_F$ as increased seed production boosts the flow of plant density through all stages. By increasing the number of plant individuals maturing, we observe a greater number of plant individuals replacing those lost to consumption. In our analysis, $\gamma_{12}$ and $\gamma_{2F}$ (along with L:D ratio) are treated as explanatory variables which we label "ecological factors," to differentiate them from other statistical inputs used in our other analyses.

**Numbered References**

39. COMPADRE Plant Matrix Database. Available from: <www.compadre-db.org>, accessed 10 August 2021, ver. 6.20.9.0. (2021).

40. Tilman, D. et al. Productivity and sustainability influenced by biodiversity in grassland ecosystems. – **Nature** *379*, 718–720 (1996).

41. Janzen, D. H. Herbivores and the number of tree species in tropical forests. – **Am. Nat.** *104*, 501–528 (1970).

42. Connell, J. H. On the role of natural enemies in preventing competitive exclusion in some marine animals and in rain forest trees. – In: Boer, P. J. and Gradwell, G. R. (eds), Dynamics of populations: proceedings of the advanced study institute on 'dynamics of numbers in populations'. Pudoc Oosterbeek, the Netherlands, 298–312 (1971).

43. Breiman, L. Statistical Modeling: The Two Cultures (with comments and a rejoinder by the author). *Statistical Science* **16**, 199–231. https ://doi.org/10.1214/ss/10092 13726 (2001b).

44. Fisher, A., Rudin, C., & Dominici, F. All models are wrong but many are useful: Variable importance for black-box, proprietary, or misspecified prediction models, using model class reliance. ArXiv e-prints. (2018).
